# CEDAR, an online resource for the reporting and exploration of complexome profiling data

**DOI:** 10.1101/2020.12.11.421172

**Authors:** Joeri van Strien, Alexander Haupt, Uwe Schulte, Hans-Peter Braun, Alfredo Cabrera-Orefice, Jyoti S. Choudhary, Felix Evers, Erika Fernandez-Vizarra, Sergio Guerrero-Castillo, Taco W.A. Kooij, Petra Páleníková, Mercedes Pardo, Cristina Ugalde, Ilka Wittig, Lars Wöhlbrand, Ulrich Brandt, Susanne Arnold, Martijn A. Huynen

**Affiliations:** Center for Molecular and Biomolecular Informatics, Radboud Institute for Molecular Life Sciences, Radboud University Medical Center, Nijmegen, The Netherlands; Institute of Physiology, Faculty of Medicine, University of Freiburg, 79104 Freiburg, Germany; Center for Biological Signalling Studies (BIOSS) and Center for Integrative Signalling Studies (CIBSS), 79104 Freiburg, Germany; Institute of Plant Genetics, Leibniz Universität Hannover, Herrenhäuser Str. 2, 30419 Hannover, Germany; Functional Proteomics, The Institute of Cancer Research, London SW7 3RP, UK; Medical Microbiology, Radboud Institute for Molecular Life Sciences, Radboud University Medical Center, Nijmegen, The Netherlands; MRC Mitochondrial Biology Unit, University of Cambridge, UK; University Children’s Research@Kinder-UKE, University Medical Center Hamburg-Eppendorf, 20246 Hamburg, Germany; Hospital 12 de Octubre Research Institute, Madrid 28041, Spain; Centro de Investigación Biomédica en Red de Enfermedades Raras (CIBERER), U723, Madrid, Spain; Functional Proteomics, Medical School, Goethe-University, 60590 Frankfurt am Main, Germany; General and Molecular Microbiology, Institute for Chemistry and Biology of the Marine Environment (ICBM), Carl von Ossietzky University of Oldenburg, Oldenburg, Germany; Radboud Center for Mitochondrial Medicine, Radboud University Medical Center, Nijmegen

## Abstract

Complexome profiling is an emerging ‘omics approach that systematically interrogates the composition of protein complexes (the complexome) of a sample, by combining biochemical separation of native protein complexes with mass-spectrometry based quantitation proteomics. The resulting fractionation profiles hold comprehensive information on the abundance and composition of the complexome, and have a high potential for reuse by experimental and computational researchers. However, the lack of a central resource that provides access to these data, reported with adequate descriptions and an analysis tool, has limited their reuse. Therefore, we established the ComplexomE profiling DAta Resource (CEDAR, www3.cmbi.umcn.nl/cedar/), an openly accessible database for depositing and exploring mass spectrometry data from complexome profiling studies. Compatibility and reusability of the data is ensured by a standardized data and reporting format containing the “minimum information required for a complexome profiling experiment” (MIACE). The data can be accessed through a user-friendly web interface, as well as programmatically using the REST API portal. Additionally, all complexome profiles available on CEDAR can be inspected directly on the website with the profile viewer tool that allows the detection of correlated profiles and inference of potential complexes. In conclusion, CEDAR is a unique, growing and invaluable resource for the study of protein complex composition and dynamics across biological systems.

## 1. Introduction

The complete inventory of multi-protein assemblies present in a sample is referred to as its complexome. Complexome profiling is an approach to systematically interrogate the composition of the complexome, as well as the interplay of its constituents, in a biological sample by coupling biochemical fractionation to mass spectrometry-based proteomics [1–3]. Samples, typically mixtures of single proteins and protein complexes, are prepared and then separated under non-denaturing conditions using a continuous biochemical method. Next, fractions are collected and individually analyzed by quantitative mass spectrometry. A number of methods are used to natively separate protein complexes in complexome profiling, such as blue-native gel electrophoresis [4–18], size exclusion chromatography [19–22] and sucrose gradient ultracentrifugation [10,23]. The result of such an experiment, i.e. a complexome profile, contains a measure of abundance in every separated fraction for each detected protein. Proteins with similar abundance profiles; i.e. that have peaks in the same fractions, are relatively likely to be part of the same protein complex. These abundance profiles can be used as input for clustering analyses, grouping proteins that are likely to occur in the same complex [1,24]. Alternatively, these abundance profiles can be used to calculate various distance or similarity metrics between proteins, which can in turn be used to generate large scale protein-protein interaction networks [14,21,25–27].

Complexome profiling has been gaining considerable popularity as a systems-level alternative to classical target-centered antibody/affinity-based (AP-MS) proteomics due to the accuracy, comprehensiveness, and the highly informative nature of the results that can be obtained from a single experiment. Thus, it has been used to discover new protein complexes or identify new members of known complexes [1,5,8,10,28–32], to elucidate complex stoichiometry [33,34], to study protein complex assembly [6,7,16,35–39], to investigate evolutionary conservation of protein complexes [40], to better understand disease mechanisms[41–44] and to study dynamic reorganization of protein interactions [45]. Furthermore, several computational tools have been developed for the processing, analysis, visualization, interpretation, and comparative analysis of complexome profiling data [5,24,26,27,36,46,47].

In summary, complexome profiling is an emerging “omics” approach, and a powerful and versatile tool to study native protein assemblies in a wide range of biological systems. Complexome profiles contain a wealth of information on large ensembles of protein complexes, while published studies necessarily focus on a small subset of detected protein complexes. Additionally, comparative analysis of multiple complexome profiles has been shown to be feasible and has proven valuable to characterize differences between complexomes [45,46]. Data resulting from similar but more established ‘omics approaches are frequently reinterrogated, exhibiting their lasting value after initial publication. Thus, the growing number of published complexomes represent a valuable but under-explored information resource, with a high propensity for reuse.

Currently, the lack of a central resource that provides easy access to these data, reported with adequate information, has limited their reuse. Here, we present CEDAR (the Complexome Profiling Data Resource, www3.cmbi.umcn.nl/cedar/), a web-accessible database for the storage, sharing and inspection of complexome profiles. The establishment of standards for the reporting of results in the form of ‘minimum information’ documents has proven effective in increasing the reuse of data from other omics approaches [48–50]. To improve the reporting of complexome profiling data, we defined a standard for the formatting and a “minimum set of information required for a complexome profiling experiment”, termed MIACE. CEDAR stores the fractionation profiles resulting from complexome profiling experiments along with MIACE-compliant documentation, provides easy access to these data and allows inspection of profiles on the web through an integrated visualization tool. Together, CEDAR and MIACE, in parallel to standardization of experimental methods [51], provide the standards and infrastructure to increase the value and accessibility of published complexome profiling studies.

## 2. Results

### 2.1. MIACE, Minimum Information About a Complexome profiling Experiment

We developed MIACE with the goal to increase the value of the growing number of complexome profiling experiments being performed. To enable reuse of the results from these experiments, the information provided needs to be sufficiently complete to allow for efficient interpretation and evaluation. The authors represent researchers from nine different research institutes, with expertise in developing the complexome profiling technique, performing complexome experiments as well as analyzing and interpreting their results to generate new biological insight. Together we reached a consensus on the minimum information to adequately describe a complexome profiling experiment, while keeping the burden for reporting at an acceptable level. A complexome profiling experiment combines multiple methods to interrogate the complexome. Some of these methods have already been addressed, at least in part, in modules of MIAPE, the minimum information about a proteomics experiment [50]. However, several aspects are unique to a complexome profiling experiment. The purpose of MIACE is to provide a complete description of all the information required for comprehensively reporting a complexome profiling experiment, for which it refers to existing documentation where applicable. MIACE consists of seven components, each describing a set of core parameters required for the description and allowing the interpretation of a complexome profiling experiment (Figure 1A). The complete MIACE specification is available as a supplementary document (Suppl. A).

**Figure 1.**
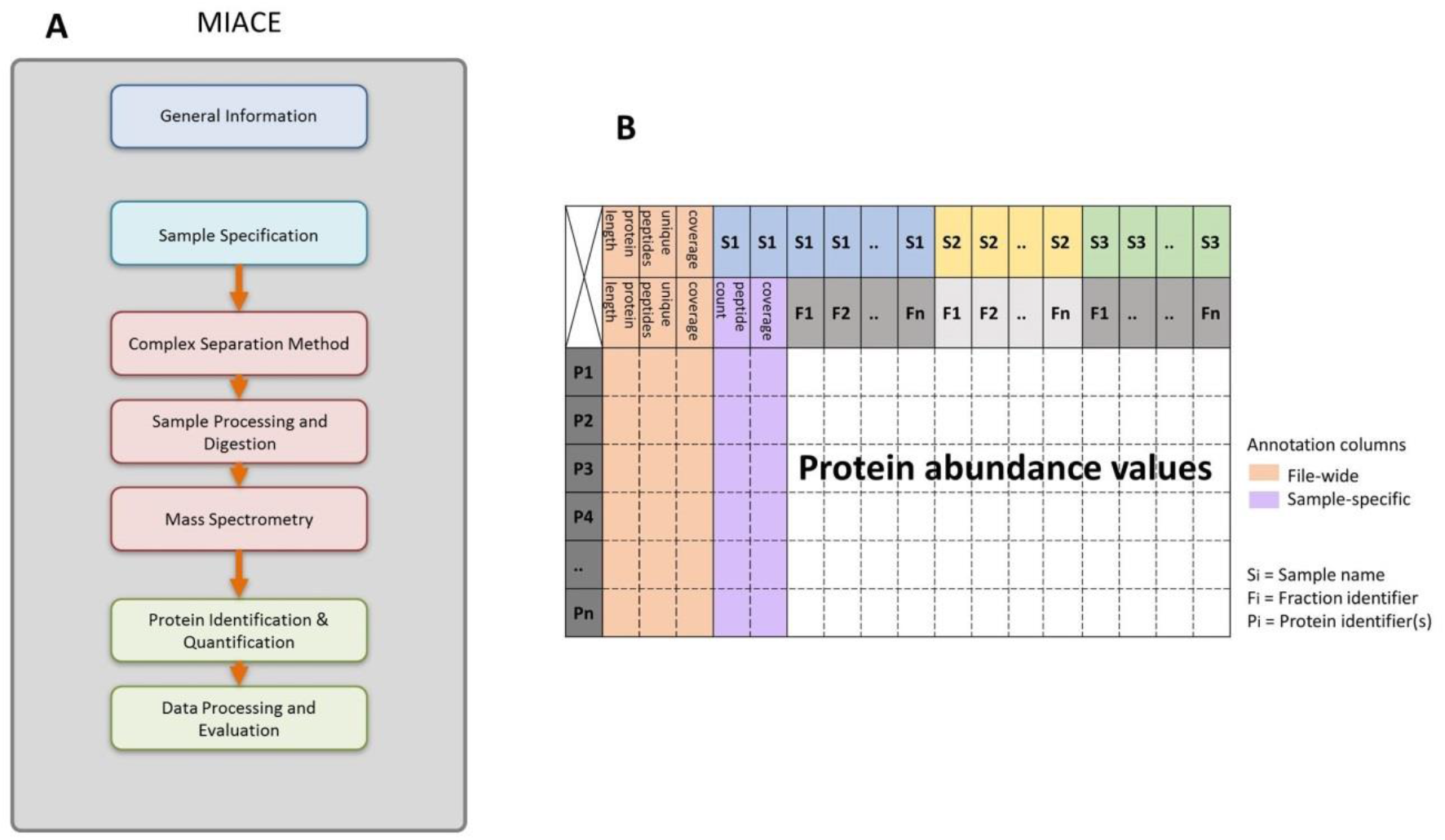
The Minimum Information About a Complexome profiling Experiment (MIACE) and the CompTab file format. A, Schematic of the 7 components of MIACE, corresponding to the steps of a complexome profiling experiment. Each component specifies the minimum information required for its adequate description. B, Structure and content of the CompTab file format, for the efficient exchange of complexome profiling data. The tabular plain text format stores the primary protein abundances per fraction in a manner that is easy to interpret by both software and researchers. In addition to the protein abundances, additional information about the protein identifications is stored.

In addition to defining requirements for the annotations and metadata of a complexome profiling experiment and as part of MIACE, we introduce a standardized file format: the complexome profiling tabular file format (CompTab) (Figure 1B). This file format is meant to store and exchange the primary results of a complexome profiling experiment, the distribution of the protein abundance data (i.e. a complexome profile). Standardizing the format for the exchange of complexome profiles will simplify sharing and reuse of these data, as well as allowing development of software that can interact with it. We chose a tabular plain text format, with the aim to be clear and simple, yet explicit, so the data can be interpreted and created by researchers and software alike. Tabular file formats have proven more practical and intuitive than others, like xml-based formats (Rayner et al., 2006). The CompTab file format specification is part of MIACE and is available as a supplementary document (Suppl. A).

### 2.2. CEDAR: Design and implementation

CEDAR has been created to make published complexome profiling data more FAIR (Findable, Accessible, Interoperable, Reusable) [52]. To achieve this, the following principles were considered in its design. Firstly, the available complexome profiles and any associated information should be easy to access. Secondly, researchers should be able to readily find complexome profiles that best meet their requirements among the available datasets. Lastly, all information required for the interpretation and reuse of the complexome profiles should be available alongside the data. **Figure 2** shows the overall structure of CEDAR. Complexome profiling data and associated information is stored in a relational database. Where feasible, aspects of the data are described using a pre-defined controlled vocabulary. This allows for efficient searching and filtering of complexome profiling data based on any associated information. Associated files are stored in file storage and linked to the relevant experiments in the database. A REST API service provides access to the database and file storage. The CEDAR website, as well as the profile viewer tool, uses this API to access the data. CEDAR provides user access to the data from the user-friendly webpage as well as direct programmatic access to the API endpoints. CEDAR makes use of the BioDBnet db2db resource [53] to retrieve additional annotations for the proteins detected in submitted complexome profiling experiments, which are then made available in the database. CEDAR is available on the web at: www3.cmbi.umcn.nl/cedar/.

**Figure 2.**
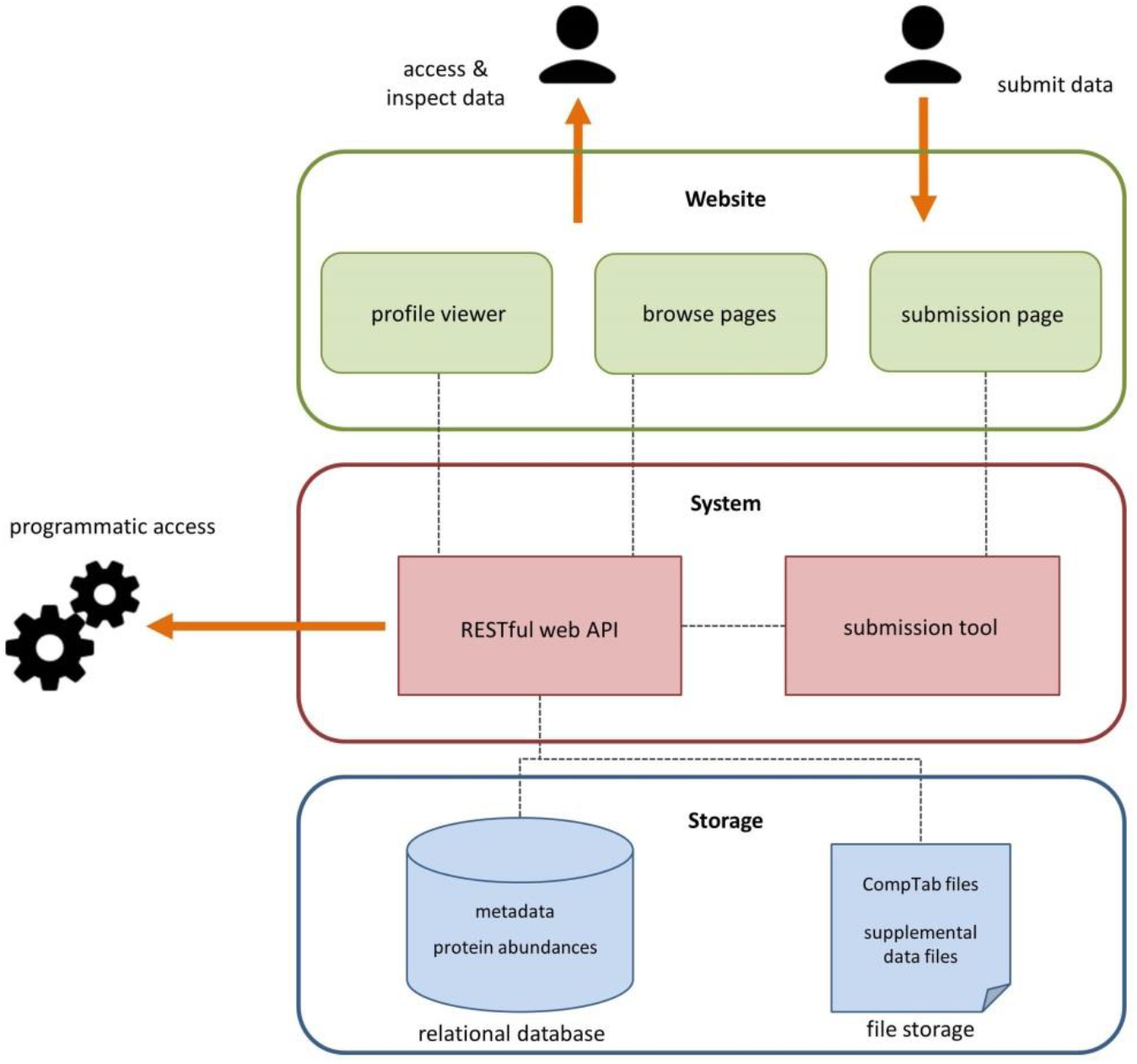
Schematic of the ComplexomE profiling Data Resource (CEDAR) structure. Experiment descriptions and complexome profiling data are stored in a relational database. Associated files are stored separately. The RESTful web interface provides access to the database and file storage. CEDAR can be accessed through a user-friendly website or programmatically by directly interacting with the restful web service. Complexome profiles can be visualized with the profile viewer web tool. The submission tool and webpage facilitate deposition of additional complexome profiling data to CEDAR.

### 2.3. CEDAR: Browsing experiments and samples

CEDAR provides access to complexome profiling data at two levels. In the experiment section one can browse through complete experiments. An experiment in CEDAR describes a related set of complexome profiling samples that are provided in a single submission, including all associated information. Each experiment has its own page, which contains detailed information about the experiment, all samples that comprise the experiment, and any associated files. Additionally, one can browse through the total collection of samples (i.e: complexome profiles) in the sample section, which is useful when one is interested in the complexome of a specific condition or system rather than the complete set of samples belonging to a certain experiment. Like experiments, each sample has its own page with detailed information, links and associated files. The search and filter functionality in the browse sections make it easy to find relevant complexome profiling data (**Figure 3**). The search function provides free-text search functionality, covering all information associated to the experiment or sample. The filter option allows filtering the experiments or samples based on specific descriptors, which are deposited using a controlled vocabulary. Additionally, for researchers that have a specific protein of interest, using the protein filter function one can select experiments or samples in which a certain protein was detected. Any information associated with an experiment can be used to find its samples, and any information associated with a sample can be used to find the corresponding experiment. In addition to experiments and samples, CEDAR stores information about which samples originate from the same complex separation run. If an experiment contains sample sets comprising multiple separate complex separation runs, information regarding the batch that the samples originate from is relevant, as there tend to be batch effects between samples from different complex separation runs [46]. Once a sample or experiment of interest has been selected, the complexome profiles can be accessed in multiple ways. The data can be downloaded as CompTab format files, which have been provided during submission. Alternatively, the abundance profiles of a specific sample, and optionally a subset of proteins, can be downloaded directly from CEDAR. This method has the option to include protein annotations provided by CEDAR alongside the primary protein accessions and the abundance data. Downloaded data can then be used in external data evaluation or further analysis, e.g. using NOVA[24], EPIC[27], PrInCE[26], ComplexomeMap[5],or other available tools.

**Figure 3.**
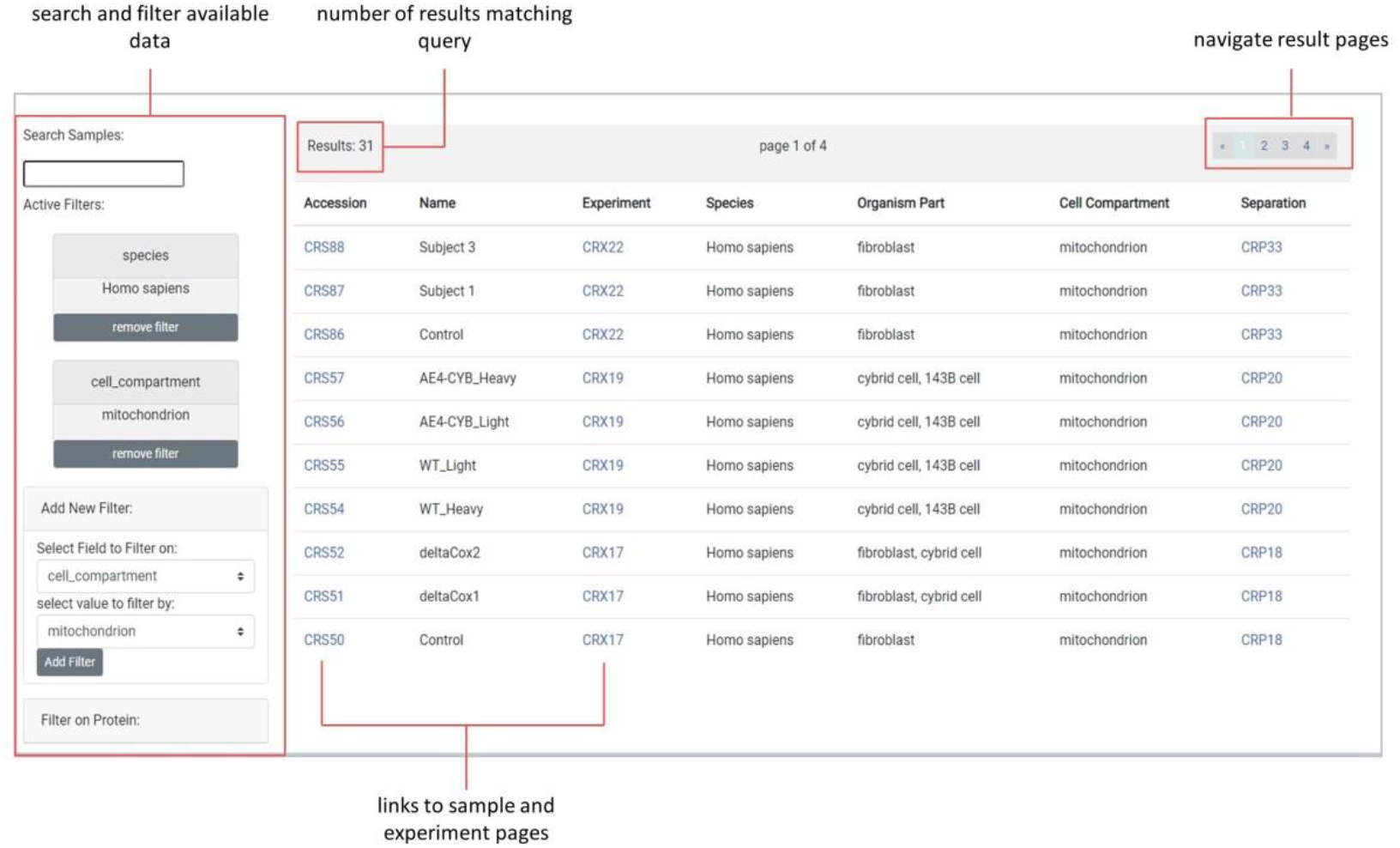
View of the CEDAR browse samples section, showing the general layout and search and filter functionality of both the sample and the experiment browse sections. The left column of the webpage has search and filter options to query for samples or experiments of interest. The general search bar allows for free text search of any experiment and sample descriptions. Filters can be added to select samples based on specific aspects, which are defined with controlled vocabularies. A protein filter can be applied to select only samples in which this protein is detected. The main table shows all samples matching the current query, along with general information and links to the relevant sample and experiment pages.

### 2.4. Interactive exploration of complexome profiles with the Profile Viewer tool

Aside from downloading the data to inspect or use it locally, inspection of any complexome profiling sample can be done directly on the website using the Profile Viewer (**Figure 4**). When viewing a sample in the Profile Viewer, the complete set of detected proteins, along with annotations and additional information, are shown in a table. The abundance profiles of one or more selected proteins are visualized in the interactive plot, with various customization options like normalisation or smoothing of the profile and zooming. In addition to visualization of selected proteins, the Profile Viewer allows on-the-fly correlation of protein abundance profiles within user-defined windows of interest, allowing inference of potential protein complexes. The profile viewer allows the calculation of Pearson correlation scores between the protein of interest and all other proteins in the dataset. The profiles of the top-scoring proteins will be also be shown in the profile plot.

**Figure 4.**
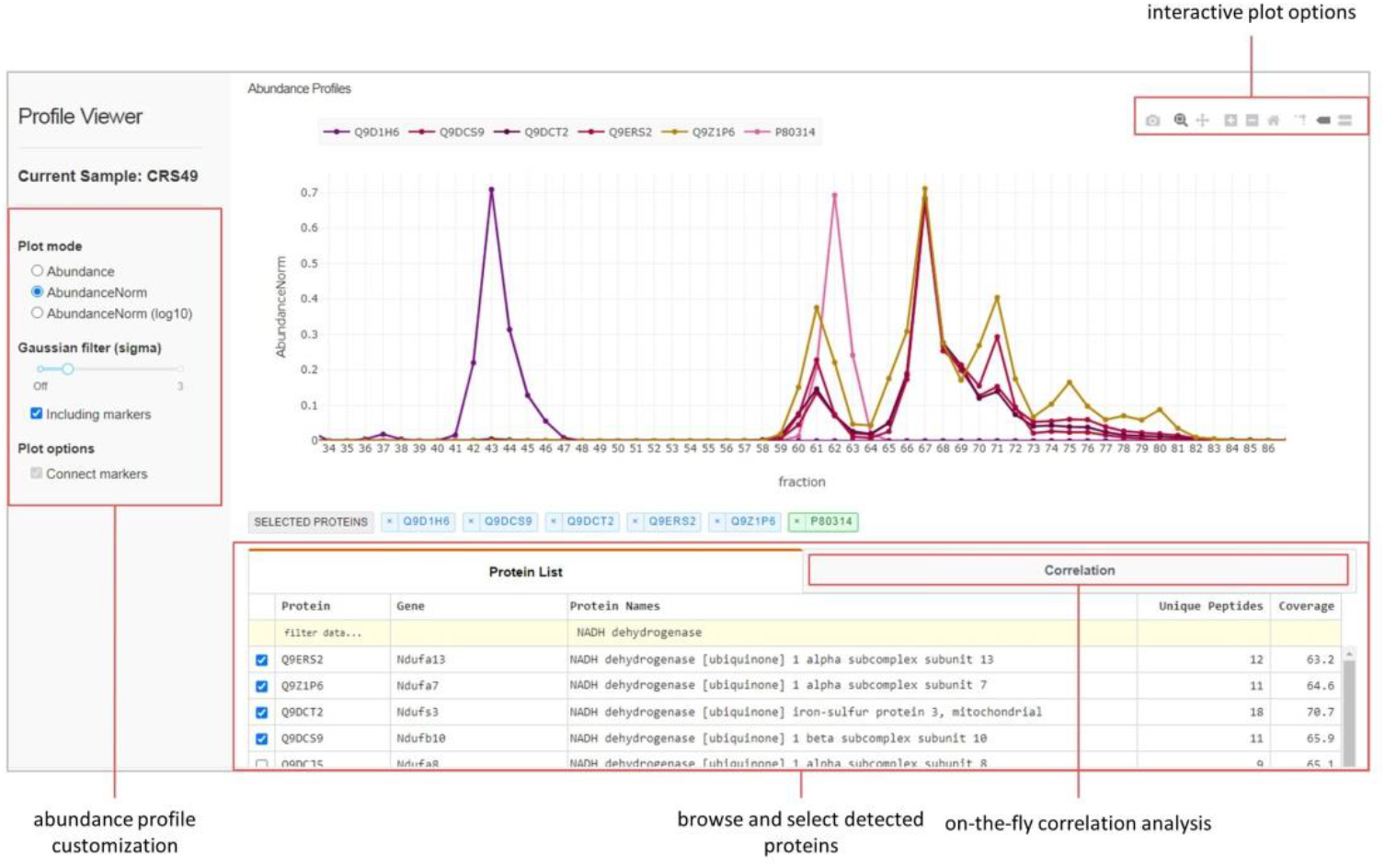
Overview of the profile viewer tool. Protein abundance profiles of any complexome profiling sample available on CEDAR can be visualized here. The protein list tab contains a table showing the complete set of proteins detected in this sample, with additional information. The interactive plot shows abundance profiles of selected proteins, in this case of NADH:ubiquinone oxidoreductase, with several options for customization. The Correlation tab allows on-the-fly correlation analysis of proteins of interest.

### 2.5. Programmatic access through CEDAR’s REST API

The RESTful web service of CEDAR provides programmatic access to the data in CEDAR. The API provides access points that can be used to query and retrieve multiple complexome profiling datasets available in CEDAR, with the same search and filter functionality as the web interface. Additionally, individual samples and experiments, including all associated data and files, can be accessed using their respective ID’s. Data can be accessed through HTTP requests, which return JSON format responses, a widely used format for the exchange of data. Documentation on how to make use of the REST service is available on the CEDAR webpage. In addition to JSON format responses CEDAR provides webpage responses for the main API endpoints, simplifying development of scripts and software that interact with the API (**Figure 5A**). These html pages, which can be viewed with a browser, provide a quick and clear overview of all the querying options and syntax, and generate URLs that would yield the result of any query manually entered on the html webpage.

**Figure 5.**
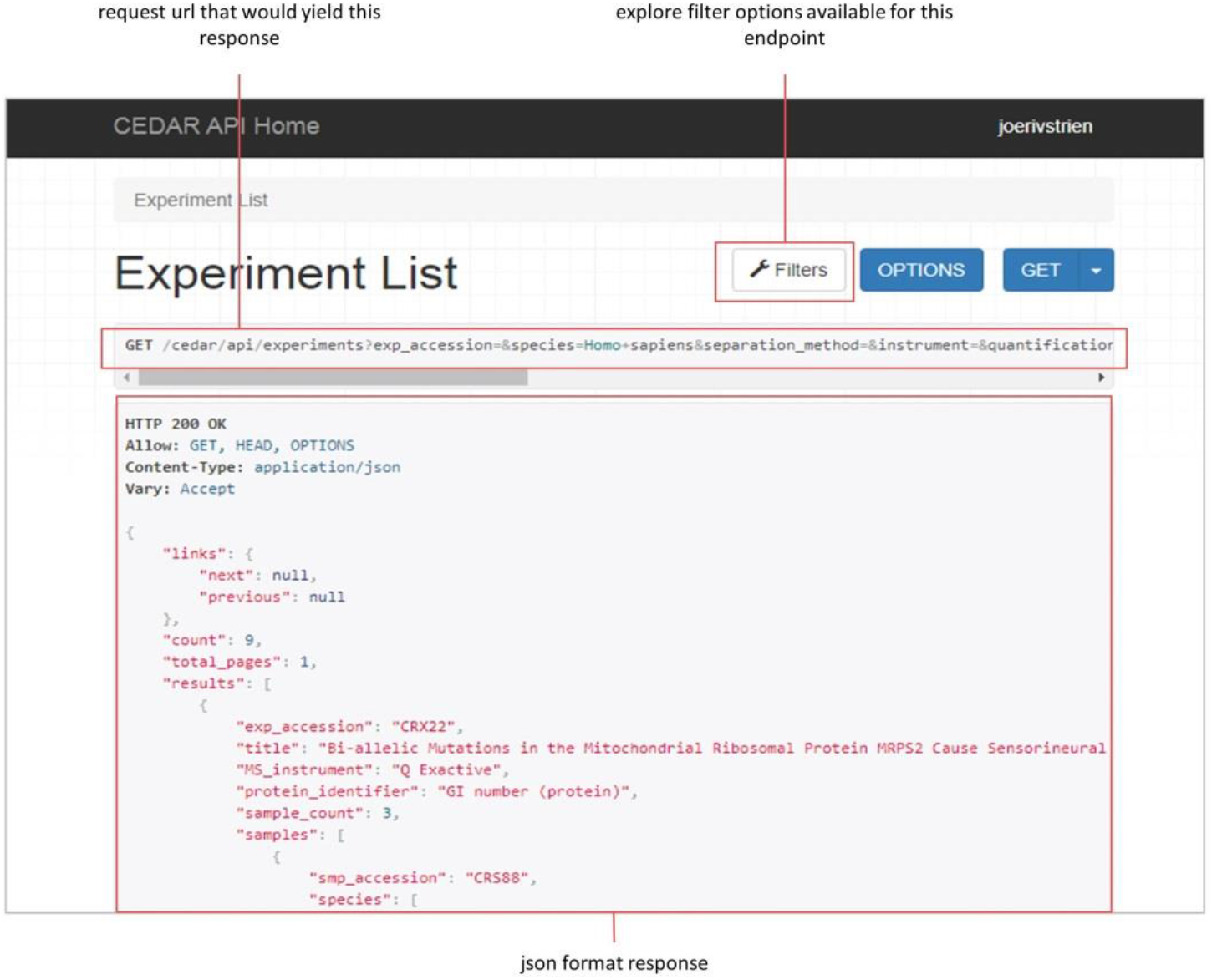
A, view of the ‘browsable’ web page for the REST API endpoint for batch retrieval of experiments. The ‘filters’ button allows exploration of available search and filter options. After entering search or filter terms, the request URL that would yield the results of this query is shown at the top. The main body of the page shows the JSON format response data for this request.

### 2.6. Submitting to CEDAR

Researchers can upload the results of their complexome profiling experiments to CEDAR using the online submission tool, which is available on the CEDAR webpage. This tool allows submission of a complete MIACE-compliant complexome profiling experiment, including data files and all required information, in a stepwise submission process. The submission tool has extensive documentation, provides feedback during the submission, and helps keeping track of the submission process, rendering a complete MIACE-compliant submission manageable. A number of descriptors in the database are defined by a controlled vocabulary, to improve querying and interoperability of data. The CEDAR submission tool interacts with the ontology lookup service [54] to provide up-to-date access to the appropriate ontologies when submitting an experiment (**Figure 5B**). To further consolidate the terminology used in the database, CEDAR initially suggests as selectable options only terms that are already used in the database as choices. If none of these terms is adequate, the full ontology can be browsed to select a term. When an experiment is initially submitted it is not publicly available, and will only be published to the website after the submission has been validated.

### 2.7. CEDAR: Available data

Results from published complexome profiling experiments have already been uploaded by their authors to CEDAR using the submission tool. While more data is being submitted regularly, we provide a snapshot of the collection of data that is currently available in CEDAR. Results from sixteen different studies, comprising 83 complexome profiling samples have been deposited, originating from a variety of biological systems: mammals (human, mouse, rat), Apicomplexa (three different *Plasmodium* species), as well as a plant and an archaeal species (*Arabidopsis thaliana* and *Candidatus* “Methanoperedens”, respectively). An overview is provided in Table 1. The majority of current complexome profiles refer to samples of isolated mitochondria from the respective organism. Additionally, CEDAR contains complexome profiles of the whole cell, or subcellular components other than mitochondria (e.g. endosome, lysosomal membrane, secretory granule). The number of proteins detected per complexome profile ranges from 709 to 4160, with an average of about 2000. The number of fractions in which the samples are separated before MS identification and quantification of proteins ranges from 23 to 231, with an average of about 60.

**Table 1.** An overview of the collection of complexome profiling data currently available on CEDAR. The uploaded experiments have been categorized by study type, the subject species, as well as by the cell compartment(s) that were studied.

| study type | species | number of experiments | number of samples | cell compartment(s) |
| --- | --- | --- | --- | --- |
| characterization of complexome | *Homo sapiens* | 1 | 4 | mitochondrion |
|  | *Rattus norvegicus* | 1 | 1 | mitochondrion |
|  | *Arabidopsis thaliana* | 1 | 1 | mitochondrion |
|  | *Plasmodium* species (*falciparum*, *knowlesi*, *berghei*) | 2 | 29 | whole cell (18), mitochondrion (11) |
|  | *Candidatus* Methanoperedens | 1 | 10 | whole cell |
| characterization of patient complexome | *Homo sapiens* | 4 | 16 | mitochondrion (10), other (6) |
| characterization of gene knockout complexome | *Homo sapiens* | 2 | 10 | mitochondrion |
|  | *Mus musculus* | 1 | 4 | mitochondrion |
| characterization of mutant cell line complexome | *Homo sapiens* | 2 | 7 | mitochondrion |
| characterization of gene knock-in complexome | *Mus musculus* | 1 | 1 | mitochondrion |

## 3. Discussion

Here we present CEDAR, an online data resource for the storage and sharing of complexome profiling data, and MIACE, a specification for the reporting of such data. The MIACE specification document represents a consensus of a number of complexome profiling experts from different institutes. Yet, these standards are open to further development, as additional insights might arise from future studies. While the MIACE standard is developed in parallel with and presented alongside CEDAR, it is an independent standard that does not intend to impose any resource or approach, but should be applicable to the reporting of any kind of complexome profiling dataset.

There are other databases that can store the results from proteomics/mass spectrometry experiments, such as those that are part of proteomeXchange [55–57]. However, they are not tailored towards efficient reuse of complexome profiling data. First, they do not provide easy access to the results (i.e. protein abundance profiles) in a consistent format. Second, these databases rather function as raw data repositories offering little support for visualization, customization or analysis of results. This complicates efficient interpretation and reuse of published complexome profiles. CEDAR stores standardized descriptions that are specific to a complexome profiling experiment, simplifying comparative analysis and independent evaluation of complexome profiles. The Profile Viewer tool leverages the availability of complexome profiles in the CEDAR database and allows for direct and interactive inspection of any available dataset. This facilitates inspection of abundance profiles to study protein complex composition and behavior, without requiring additional software or experience working with these data. This should lower the threshold for the broader scientific community to make use of the available complexomes. CEDAR stores the final results of a complexome profiling experiment. The original mass spectrometry output is however a valuable resource, for example for the reanalysis using different tools or identification/quantification methods, and can and should still be deposited in databanks like those part of proteomeXchange. An experiment submitted to CEDAR can be linked to these data using a proteomeXchange identifier. The REST API portal allows for automated access to data in CEDAR by software tools, and allows interaction with CEDAR from any programming or scripting language. This facilitates data mining and large-scale reanalysis of complexome profiles, as well as the development of further software for the analysis and visualization of these data.

CEDAR presents an accessible and rich information resource that will continue to grow as more existing and still to be generated complexome profiling data are submitted to CEDAR. The facile access of complexome profiling data will help answer new questions regarding the differences in the complexomes between conditions and systems or aspects of the complexome that have not been addressed yet. As a result, insights beyond what was uncovered in the original publication of these data can be expected. For researchers generating complexome profiling data, comparison of their findings with available complexome profiles will help place their findings into a broader context. Additionally, large-scale analysis leveraging many complexome profiles will help achieve the sensitivity necessary to study elements of complexomes that have remained hidden so far. The proper standard and infrastructure provided will push the complexome profiling method forward, and empower researchers to better understand the roles of protein complexes and interactions in the functioning and malfunctioning of a wide variety of systems. We aim to make submission of complexome profiling data to CEDAR a standard upon acceptance of manuscripts.

## Acknowledgements

This work was supported by grants from the Dutch Organisation for Health Research and Development (ZON-MW TOP grant number 91217009; MH., UB, SA and JS), the Netherlands Organisation for Scientific Research (NWO-VIDI 864.13.009, TWAK and FE), Comunidad Autónoma de Madrid (ERDF-ESF Grant P2018/BAA-4403; CU), Instituto de Salud Carlos III-MINECO (European FEDER Funds Grant PI17-00048; CU), the German Research Foundation (DFG) under Germany’s Excellence Strategy (CIBSS – EXC-2189 – Project ID 390939984; US), DFG grant (SFB 746, TP 20; US), the Medical Research Council Core (Grant MC_UU_00015/5; EFV), the Association Française contre les Myopathies (AFM) (Grant 16086; EFV), the Deutsche Forschungsgemeinschaft (SFB 815/Z1; IW), BMBF mitoNET — German Network for Mitochondrial Disorders (01GM1906D; IW), Medical Research Council, UK (MC_UU_00015/4, PP), Marie Sklodowska-Curie ITN-REMIX grant (Grant 721757; PP), DFG grant (Br 1829/16-1; HPB), and the CRUK Centre (C309/A25144).

