## Supplement A for "CEDAR, an online resource for the reporting and exploration of complexome profiling data"

Suggested (meta-)data standards for the reporting of complexome profiling experiments

Version 1.0

* corresponding author

**A proposal of (meta-)data standards for the reporting of complexome profiling experiments**

**introduction**

Complexome profiling is an emerging ‘omics approach that interrogates protein complex composition, assembly and dynamics [1,2,11–20,3,21,4–10]. This approach combines biochemical methods to separate protein complexes present in a biological sample into a series of fractions, with mass spectrometry to determine the protein content in each of the separated fractions, to then combine this information and create a protein distribution profile.  The generated data inherently contains comprehensive information on a large collection of multiprotein assemblies present in the studied system. It should be noted that the published complexome profiling studies have generally focused on a small subset of the protein complexes detected in the collected datasets. Thus, the potential for reusability of these data is very high, e.g. to uncover the comigration of proteins with each other in independent experiments. Nevertheless, a standard for its reporting, storing and annotation has not yet been defined, limiting the optimal use of this valuable and growing information resource.

The establishment of standards for reporting of a certain experiment through a minimum information document has proven to be very effective, greatly improving the availability and (re-)usability of the results of, for instance, microarray experiments, high-throughput sequencing, and various proteomics approaches [22–24]. Therefore, we propose the MIACE standard, the Minimum Information About a Complexome profiling Experiment. MIACE is created in order to facilitate efficient interpretation, sharing and reuse of complexome profiling data. It consists of the following seven components, each describing a set of core parameters required for the description and interpretation of a complexome profiling experiment:

1. General information
2. Sample specification
3. Protein separation method(s)
4. Sample processing and digestion
5. Mass spectrometry
6. Protein identification and quantification
7. Data processing and evaluation

Several aspects of these components are unique to a complexome profiling experiment, i.e. the separation of the native complexes into discrete fractions, and the calibration of apparent complex mass corresponding to these fractions with the help of known soluble and/or membrane bound reference complexes. Others (protein separation, mass spectrometry, protein identification/quantification and data processing/evaluation) are at least in part already addressed in modules of MIAPE, the Minimum Information About a Proteomics Experiment (www.psidev.info/miape)[22]. Where appropriate, we refer to these MIAPE components, such as gel electrophoresis, column chromatography, mass spectrometry, mass spectrometry informatics[25–28], for a description of the minimum information required. This document combines all information required for reporting a complete complexome profiling experiment.

**1. General information**

This section contains a general description of the experiment, which is the complete set of related complexome profiling samples that together address a certain scientific question. An experiment typically (but not necessarily) refers to the complete collection of complexome profiling data published in a single paper. It includes:

a) information about the author, affiliation(s), contact information.
b) short, single sentence title of the experiment.
c) longer general description of the experimental design and setup.
d) links to other related resources, reference to a publication if applicable.

e) relationship between all samples and associated data files as well as any information about relationships between samples. For example, which samples originate from the same batch or gel. This is best represented as a table, containing all samples, data and their relations.

### 2. Sample specification

This section contains a detailed description of all samples comprising the experiment. It should include:

a) the species origin(s) of the sample.
b) the strain, cell type and/or tissue type.
c) if applicable, the specific cell compartment or type of protein complex that has been isolated or for which the sample was enriched.
d) any experimental /parameters, which could for example be a treatment, a (disease) condition.
e) if applicable, any isotope label/mass tag or other chemical derivatization performed.
f) any additional information that is deemed relevant by the author/submitter, like for example sex, age or developmental stage.

### 3. Protein separation method(s)

This section should contain a description of the method used for separating protein complexes under native conditions prior to protein identification by mass spectrometry. This can be achieved by various methods, and new methods will likely be developed in the future. These methods include, but are not limited to, separation by gel electrophoresis, size exclusion chromatography, field-flow fractionation and sucrose density gradient ultracentrifugation. Detailed explanations on the information required for the description of gel electrophoresis and column chromatography are present in the respective MIAPE components (www.psidev.info/miape) [25,26]. In general, a description of the protein separation method should include the following:

a) a description of the specific method(s) of protein separation.
b) a description of or reference to the protocol(s) used.
c) the specific conditions or parameters under which the separation took place.
d) the number of fractions into which the protein complexes are separated, and if applicable, the molecular mass range that is covered by these fractions. This can be specified per sample if samples have varying numbers of fractions or mass ranges.
e) any other information deemed necessary to accurately and fully describe the method of protein separation.
f) if technically feasible, a visualization of the separation result, like an image of the resulting gel with visible bands in the case of native gel electrophoresis, or UV traces in the case of chromatography.

### 4. Sample processing and digestion

This section should contain a description of any additional processing steps the samples have undergone, along with a description of the digestion process to prepare samples for mass spectrometry. This section should include:

a) the protease(s) used for digestion.
b) description of or reference to the digestion protocol used.
c) description of or reference to protocols of any other processing steps or treatments that were performed on the samples prior to the mass spectrometry analysis.

**5. Mass spectrometry**

This section contains descriptions of the instruments and settings used for liquid chromatography (LC) coupled with mass spectrometry (MS). For a complete description of the information required to report a mass spectrometry experiment, refer to the column chromatography and mass spectrometry components of MIAPE (www.psidev.info/miape)[26,29]. The main aspects are briefly restated here:

a) chromatography column and LC instrument details.
b) MS instrument type and details.
c) any customizations or parameter settings of the LC-MS instruments.
d) description of the ion source, and the relevant parameters.
e) a description of the analyzer, the type and its relevant parameters.

**6. Protein identification and quantification**

This section contains the mass spectrometry output, the processing of primary (i.e. LC-MS and MS/MS raw) data, and peptide and protein assignment and abundance determination in each sample. For a complete description of the information required to report the output of an analyzed mass spectrometry experiment, refer to the mass spectrometry informatics component of MIAPE (www.psidev.info/miape)[28]. The main aspects are briefly restated here:

a) if feasible, (a link to) 'raw' mass spectrometry output data, to enable re-analysis with different tools/algorithms.
b) peptide and protein identification including the used software, search database and parameters, allowed modifications, mass calibration, m/z tolerance, scores and error estimates (FDR).
c) peptide-level abundance determination (software for feature extraction, m/z and retention time alignment and assignment to peptides).
d) protein abundance determination including a description of the processing.
e) if applicable, the final data after any additional processing steps.
f) the quantification metric in which protein abundances are expressed, for example iBAQ, LFQ or intensity.

To these parameters we add a proposed file format for the storage of protein abundances of a complexome profiling experiment. The protein abundance data should be made available in the 'CompTab' format, which is specified in the associated MIACE CompTab document. These data should provide the original protein abundances, before normalization or alignment, and represent the protein abundances from a single replicate. The table should include the protein length, the number of identified peptides and the peptide coverage of the protein. Additionally, the apparent protein complex mass range for each fraction should be indicated, if applicable.

### 7. Data processing and evaluation

This section contains the description of any additional data processing steps performed on the data after computation of protein abundances, such as the clustering method used to reconstruct protein complexes, as well as any normalization, alignment, mass calibration etc. Additionally, this section contains a description of the quality evaluation of the experimental results. This should include the following:

a) tools and algorithms used for post-processing steps, with versions and all relevant settings and parameters.
b) the reference proteins/complexes and regression used for the apparent molecular mass calibration.
c) if available, any measures of the resolution of complex separation, as well as measures of quality of the protein identification and quantitation (false positive and negative assignment rates, dynamic range of abundance etc.).

### Intended use

The MIACE standard is meant to be used as a minimal descriptor of a complexome profiling experiment. Authors are encouraged to supply additional information as they deem necessary for a full description of the reported experiment. Data archives should include the seven components of MIACE as a requirement, but allow for reporting of additional information. Use of controlled vocabulary in the annotation of experiments will further improve the potential and ease of data reuse. An additional goal of this document is to prompt the development of software and data archives compliant with MIACE, which will further facilitate adoption of this standard. Additionally, this minimum information could be required and checked by journals and reviewers when complexome profiling experiments are reported or published.

### Conclusion

Community-wide adoption of this standard will greatly benefit research using and reusing complexome profiling data. It will make interpretation and verification of analysis results easier, and will enable large-scale meta-analysis and reuse of (published) complexome profiling data to better quantify the association between proteins into complexes.

### The CompTab file format specification

We propose CompTab (COMPlexome profiling TABular data format), a data format for the storage of protein abundance measurements of complexome profiling data. It is meant to be used alongside MIACE, with the goal to standardize the format in which complexome profiling data is shared and stored. It aims to be a clear and simple, yet explicit format that can be used and created by researchers and software alike. The proposed data format stores protein abundance values, as well as the following information about the identified proteins: protein length, number of unique peptides assigned to the protein and peptide coverage of the protein.

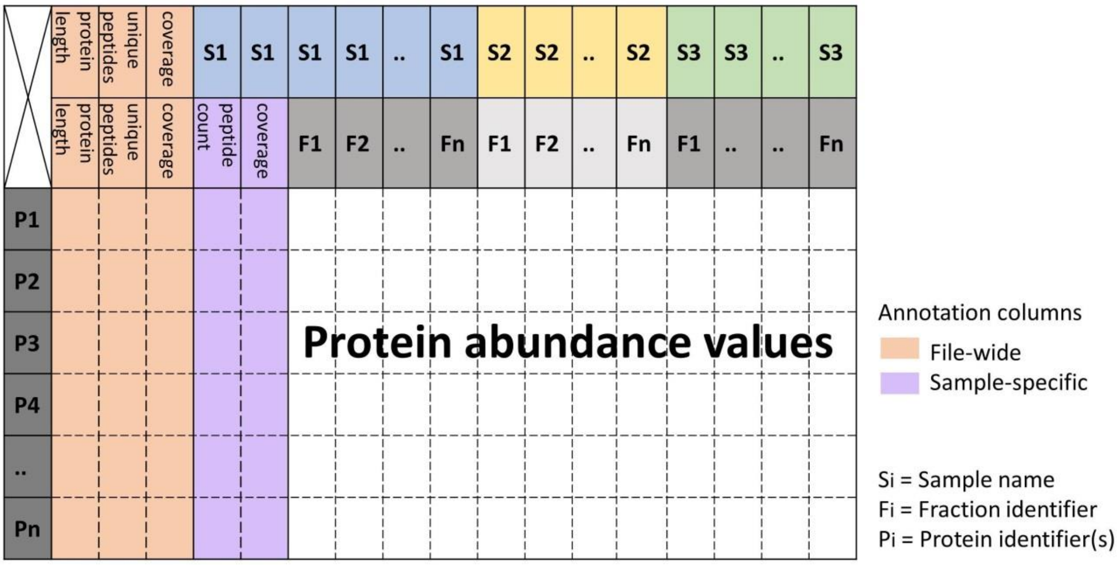


Figure 1. schematic representation of the CompTab format data structure.

#### File structure

The data is stored as a plain text file, with tab-separated values. Two header rows at the top specify the sample as well as the fraction identifiers. The first column contains protein identifiers. Each row can contain a single identifier, or in the case of protein groups, multiple comma-separated protein identifiers. If the sample fractions separate the protein complexes on their mass, the fractions are to be ordered so that lowest mass is in the leftmost column, and the heaviest fraction is in the rightmost. The columns with protein abundance profiles should contain numeric values only. In addition to the columns containing protein abundances, three types of additional columns are required, containing information about the identified proteins. The 'coverage' column provides the percent of the protein sequence that is covered by peptides. Values in the coverage column should be numeric only, without characters like ' %'.The 'unique peptides' column contains the number of different peptides assigned to the protein using whole numbers. The 'protein length' column contains the amino acid length of the protein sequence, which should be presented as integers. If the top header row of these columns specifies a sample, they are associated with this specific sample. If both header rows contain the column name (i.e. 'coverage', 'unique peptides' etc.) these columns refer to all samples in the CompTab file. The position or order of these columns in the file does not matter, as long as the headers are correct.
